# The R1-weighted connectome: complementing brain networks with a myelin-sensitive measure

**DOI:** 10.1101/2020.08.06.237941

**Authors:** Tommy Boshkovski, Ljupco Kocarev, Julien Cohen-Adad, Bratislav Mišić, Stéphane Lehéricy, Nikola Stikov, Matteo Mancini

**Affiliations:** NeuroPoly Lab, Polytechnique Montreal, Montreal, Canada; Macedonian Academy of Sciences and Arts, Skopje, Macedonia; Department of Neurosciences, Faculty of Medicine, University of Montreal, Montreal, QC, Canada; Functional Neuroimaging Unit, Centre de recherche de l’institut universitaire de gériatrie de Montréal, Montreal, QC, Canada; Montreal Neurological Institute, Montreal, QC, Canada; Paris Brain Institute (ICM), Centre for NeuroImaging Research (CENIR), Inserm U 1127, CNRS UMR 7225, Sorbonne Université, F-75013, Paris, France; Montreal Heart Institute, Montreal, QC, Canada; Department of Neuroscience, Brighton and Sussex Medical School, University of Sussex, Brighton, United Kingdom; CUBRIC, Cardiff University, Cardiff, United Kingdom

## Abstract

Myelin plays a crucial role in how well information travels between brain regions. Many neurological diseases affect the myelin in the white matter, making myelin-sensitive metrics derived from quantitative MRI of potential interest for early detection and prognosis of those conditions. Complementing the structural connectome, obtained with diffusion MRI tractography, with a myelin sensitive measure could result in a more complete model of structural brain connectivity and give better insight into how the myeloarchitecture relates to brain function. In this work we weight the connectome by the longitudinal relaxation rate (R1) as a measure sensitive to myelin, and then we assess its added value by comparing it with connectomes weighted by the number of streamlines (NOS). Our analysis reveals differences between the two connectomes both in the distribution of their weights and the modular organization. Additionally, the rank-based analysis shows that R1 is able to separate different classes (unimodal and transmodal), following a functional gradient. Overall, the R1-weighted connectome provides a different perspective on structural connectivity taking into account white matter myeloarchitecture.

**Author summary:** In the present work, we integrate a myelin sensitive MRI metric into the connectome and compare it with a connectome weighted with a standard diffusion-derived metric, number of streamlines (NOS). Our analysis shows that the R1-weighted connectome complements the NOS-weighted connectome. We show that the R1-weighted average distribution does not follow the same trend as the NOS strength distribution, and the two connectomes exhibit different modular organization. We also show that unimodal cortical regions tend to be connected by more streamlines, but the connections exhibit a lower R1-weighted average, while the transmodal regions tend to have a higher R1-weighted average but fewer streamlines. In terms of network communication, this could imply that the unimodal regions require more connections with lower myelination, whereas the transmodal regions take more myelinated, but fewer, connections for a reliable transfer of information.

## Introduction

The brain is a complex system that can be modelled as an intricate network of interconnected elements (Fornito et al., 2016). Using magnetic resonance imaging (MRI), connectomics strives to characterize macroscopic connectivity by viewing the brain as a set of nodes defined by functionally or anatomically distinguishable regions of interest (ROIs) and edges that are conventionally assumed to reflect the white matter tracts connecting those nodes (Bassett & Sporns, 2017; Hagmann et al., 2007; M. P. van den Heuvel et al., 2008). Specifically, the white matter tracts can be reconstructed using diffusion MRI and tractography (Jeurissen et al., 2019; Mori & Van Zijl, 2002). In addition to delineating the nodes and edges of a brain network, weights can be assigned to the connections, which are presumed to reflect relevant properties (Rubinov & Sporns, 2010).

There is an ongoing debate as to the most appropriate choice of weighting for the connectome (Yeh et al., 2020). So far, the most widely used weight is the number of streamlines (NOS), which counts the reconstructed streamlines, from diffusion tractography, between pairs of ROIs (Fornito et al., 2016). This measure theorizes that the NOS is associated with the axonal fibers between pairs of ROIs. In fact, previous work (Sinke et al., 2018; Martijn P. van den Heuvel et al., 2015) showed a positive correlation between NOS and tract-tracing connectivity, suggesting that NOS could be used in principle as a proxy for microstructural fiber count. Despite its extensive use and the mentioned relationship with tract-tracing connectivity, the use of NOS to weight the structural connectome is still problematic (Calamante, 2019). In particular, the NOS does not measure biologically meaningful properties such as conduction velocity or channel capacity. Additionally, fiber tracking often lacks specificity as it can be affected by a number of factors, including the tractography algorithm used (Derek K Jones, 2010; Yeh et al., 2020) as well as image acquisition parameters (Derek K. Jones et al., 2013).

Another potential candidate for weighting the connections is the fractional anisotropy (FA) that can be obtained using diffusion tensor imaging (DTI). While FA does provide more insights about the microstructural properties of white matter, it is influenced as well by numerous tissue properties, including axonal diameter, fiber density, tissue geometry, as well as the degree of myelination (Derek K. Jones et al., 2013). In order to close the gap between structure and function in the connectome, a different and more meaningful choice would be characterizing white matter connections by their electrical conduction properties. A complementary approach to achieve this is given by collateral quantitative magnetic resonance imaging (qMRI) techniques, which focus on measuring physical properties of biological tissues that can be used to reconstruct the underlying microstructure. Specific qMRI measures (i.e. magnetization transfer ratio [MTR], longitudinal relaxation rate [R1], myelin water fraction [MWF]) can be used to estimate an important determinant of conduction velocity, myelin. Myelin is the dielectric material that wraps around the axons to enable fast conduction in the brain. It has recently been reported that the myelination seems to be also activity-dependent, providing a potential plasticity mechanism (Sampaio-Baptista & Johansen-Berg, 2017). Moreover, the use of such metrics is particularly well-suited for studies that examine pathology related to myelin-specific changes in brain connectivity.

Several studies (Kamagata et al., 2019; Mancini et al., 2018; Martijn P van den Heuvel et al., 2010) used such myelin-sensitive MR measures in brain network models. A recent study (Mancini et al., 2018) used the g-ratio, which is defined as the ratio between the inner and outer axonal diameters, derived from MRI as a myelin weighting scheme for the connectome. Van den Heuvel and colleagues (Martijn P van den Heuvel et al., 2010) used magnetization transfer ratio (MTR) to assess network structure in schizophrenia. Similarly, Kamagata et al. (Kamagata et al., 2019) used the g-ratio to weight the connectome in multiple sclerosis (MS) patients because of its increased sensitivity to demyelination. In the work of (Caeyenberghs et al., 2016), multiple quantitative myelin-sensitive MRI metrics were used as weights, including the R1, which has been shown to be effective for myelin imaging (Stüber et al., 2014). Caeyenberghs et al. analyzed the white matter plasticity using connectomics to determine which measures best correlate with white matter plasticity during a working memory task. The R1 measure is implemented as a stock sequence on many commercial scanners (e.g. MP2RAGE), making it easy to use out of the box.

In this paper, we integrate a myelin-sensitive measure (R1) into the structural connectome. We hypothesize that the R1-weighted connectome complements the network organization of the NOS-weighted connectome. In order to evaluate this, we weighted the connections in the structural connectome using the median R1 value along a bundle of streamlines connecting pairs of brain regions. We then compared the R1-weighted connectome with the conventional NOS-weighted connectome in terms of multiple network attributes, including strength distribution and modular structure. We then looked at different cytoarchitectonic and functional classes and networks to further explore the empirical modular structure. These differences between the R1- and NOS-weighted connectomes in terms of their overall network organization have the potential to provide a complementary perspective on white matter myeloarchitecture since R1 is more directly sensitive to myelin compared to NOS that is considered as an index of microstructural fiber count.

## Materials and methods

### Data acquisition

35 healthy volunteers (HC) (12 F/ 23 M, mean age ± sd: 61.2 ± 9.16 years) participated in the present study. Subjects were scanned at the Paris Brain Institute (ICM – Institut Cerveau Moelle), Paris, France. Scans were performed on a 3T SIEMENS Prisma Scanner. The protocol included: i) 3-shell DWI sequence (TR = 10,400 ms, TE = 59 ms, voxel size=1.7×1.7×1.7 mm^3^, number of gradient directions (per shell) = 64, 32, 8 at respectively b=2500, 700, 300 s/mm^2^, s), ii) magnetization-prepared 2 rapid acquisition gradient echoes (MP2RAGE) sequence for quantitative R1 mapping (TR=5000 ms, TE = 2.98 ms, Flip Angles = 4° and 5°, TI=700/2700 ms, FOV=256×232 mm, voxel size=1 mm3).

#### Reconstruction of quantitative R1 maps

The MP2RAGE sequence (Marques et al., 2010) produces two T1-weighted images with different flip angles and different inversion times (INV1 and INV2). These images are then combined to produce a more uniform T1w image (UNI). Then, the UNI image was used to estimate the quantitative longitudinal relaxometry maps (T1 maps) using qMRLab (Cabana et al., 2015). The longitudinal relaxation rate (R1) was then calculated from the T1 maps as:

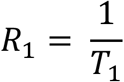

The quantitative maps were reconstructed using the qMRLab module MP2RAGE (Cabana et al., 2015).

#### Anatomical and diffusion data pre-processing

As a first step in the anatomical preprocessing pipeline, background noise removal (O’Brien et al., 2014) was applied to the UNI images using a combination of the two inversion time images with a denoising regularization factor of 70. The denoised UNI images were then processed using FreeSurfer 6.0 (Fischl, 2012) to segment the different tissues and parcellate the brain using the Desikan-Killiany Atlas (Desikan et al., 2006). In order to reduce the bias from the different parcel sizes, we subdivided them into finer regions of approximately equal size using the Lausanne 2008 parcellation (scale 125) (Cammoun et al., 2012; Hagmann et al., 2008), which resulted in 234 brain parcels. Furthermore, because this article focuses on the connectivity between cortical regions, we discarded all the subcortical regions from the analysis, which resulted in 219 brain regions.

The preprocessed anatomical images, T1w image and parcellation, as well as the reconstructed quantitative maps for each subject were transferred to the subject’s diffusion space using FSL FLIRT (Jenkinson et al., 2002, 2012) rigid body registration. The preprocessing of the diffusion images was done using MRtrix3 (Tournier et al., 2019). First, we applied a noise removal technique (Veraart, Fieremans, et al., 2016; Veraart, Novikov, et al., 2016) followed by a Gibbs ringing artifacts removal method (Kellner et al., 2016) and a B1 field inhomogeneity correction. Then, the images were preprocessed for motion and inhomogeneity distortion correction using FSL’s eddy (Andersson & Sotiropoulos, 2016) and topup tools (Andersson et al., 2003) respectively. Furthermore, in order to increase the anatomical contrast and improve the tractography and registration, the preprocessed images were upsampled to a 1mm isotropic resolution. Multi-tissue constrained spherical deconvolution (Jeurissen et al., 2014), followed by the anatomically-constrained tractography method (Smith et al., 2012), were used to reconstruct the tractogram. We applied the SD_STREAM deterministic tracking algorithm (Tournier et al., 2012) that used 1 million seeds dynamically placed using the SIFT mode (Smith et al., 2015). The tractography procedure was set to stop either when: (i) it produces 200,000 streamlines and/or (ii) the maximum number of seeds (1,000,000) is reached. During tracking the maximum turning angle was set to 60°. Streamlines with length shorter than 20 mm or longer than 250 mm were discarded from the tractogram. Additional constraints were provided by the Anatomically-constrained tractography (ACT) framework (Smith et al., 2012).

#### Structural connectome reconstruction

Structural connectivity was represented using a weighted graph, where each node corresponded to one of the 219 cortical ROIs, and each edge reflected the presence of reconstructed streamlines between each pair of ROIs. Two metrics were used as weights of the connections: (i) the NOS reconstructed between two regions and (ii) the median R1 values along the bundle of reconstructed streamlines between two regions. The same steps were followed to reconstruct the FA-weighted connectome (see Supplementary Materials). In order to mitigate the problem with spurious connections reconstructed by the tractography algorithm, we considered two nodes as connected only if there are at least 2 streamlines connecting the specific pair of ROIs. Also, a more conservative threshold (at least 5 connections) was applied to test the robustness of the results.

A group-consensus approach for both NOS- and R1-weighted connectomes was adopted to reduce individual variability in the reconstructed networks. The group consensus networks for both connectomes were constructed by taking into account only the connections that are present in at least 50% of the subjects (de Reus & van den Heuvel, 2013). The weight of a connection in the group consensus network corresponded to the median of the connection’s weights across subjects. We then assessed the relationship between the connections weights of the R1-weighted connectome and the NOS-weighted connectome using linear regression, as well as between the R1-weighted connectome and the FA-weighted connectome.

### NOS strength and R1 weighted average

We chose strength as a measure of centrality because of its straightforward interpretation. For the NOS-weighted connectome, the strength was calculated as:

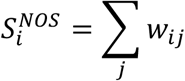

where *i* is a given node, *w*_*ij*_ is the NOS connectivity between the nodes *i* and *j*.

For the R1-weighted connectome, we looked at the R1-weighted average as it is not influenced by the number of connections (Kamagata et al., 2019). The R1-weighted average was calculated as:

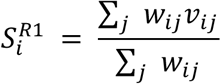

where *i* is a given node, *w*_*ij*_ is the number of streamlines and *v*_*ij*_ is the median R1 sampled along the bundle of those streamlines connecting the nodes *i* and *j*.

We then looked at the distribution of the centrality measures for each weight. The nodes were first sorted according to their NOS strength. Then, we defined the hubs as regions that have NOS strength of at least 2 standard deviations above the mean NOS strength (Martijn P. van den Heuvel & Sporns, 2013). A more conservative hub definition, at least 3 standard deviations above the mean NOS strength, was also used. Then, we highlighted the hub regions, defined in the NOS-weighted connectome, in the R1-weighted connectome.

### Modular Structure

In order to probe the modular structure of the NOS- and R1-weighted connectomes, we used a modularity maximization method (Blondel et al., 2008; Rubinov & Sporns, 2011; Sporns & Betzel, 2016). This is a common method that is used to divide a network into modules/communities with highly interconnected regions within, and less connected regions between the submodules. To achieve this, the method aims to maximize a quality function given by the following equation:

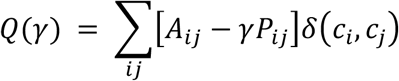

where *A* is the empirical connectivity matrix, and *P* represents the estimated connectivity matrix given a specific null model. The module assignment of node i is described by the variable *c*_*i*_, whereby *δ*(*c*_*i*_, *c*_*j*_) is the Kronecker function which is equal to 1 when *c*_*i*_ = *c*_*j*_ and 0 otherwise.

The modularity maximization also depends on a resolution parameter (γ) which makes it sensitive to different scales. If γ<1, then the network is partitioned into larger modules, while for γ>1 the method tends to find smaller modules.

To determine at which resolution the modular structure is best described, i.e. when it maximizes the quality functions, for each connectome, we iterated the method over γ values ranging from 0.5 to 3 with steps of 0.1. At each step, we ran the Louvain algorithm 1000 times (Blondel et al., 2008). Then, the resolution with highest Q was selected by taking the one that resulted with the highest Rand index (Traud et al., 2011) similarity and created a consensus modularity using the netneurotools package (https://github.com/netneurolab/netneurotools).

#### Rank-based analysis

To further explore the modular structure and assess the difference between weights, a rank-based analysis (Vázquez-Rodríguez et al., 2019) was performed: the nodes were first sorted by their strength (for the NOS-weighted connectome) and by their weighted average (for R1-weighted connectome) defining their nodal rank (1 meaning highest and 219 meaning lowest). Then, nodal ranks in the NOS-connectome were subtracted from the corresponding nodal ranks in the R1-weighted connectome. To normalize the difference, a z-score normalization was applied. The nodes were then grouped according to the von Economo cytoarchitectonic parcellation (Scholtens et al., 2018) and Yeo’s functional parcellation (Yeo et al., 2011). Finally, the median z-score for each cytoarchitectonic and functional class was computed across the respective nodes.

## Results

In order to assess the shared variance between the different connectomes, we first compared the connection weights of the R1-weighted connectome with the ones of NOS- and FA-weighted connectomes. We found that the R1 and NOS weights exhibited an R2 of 0.023 (p<0.01), while the R1 and the FA weights exhibited R2 of 0.24 (p<0.01) (Figure 1). Given that R1 measures different microstructural properties compared to NOS and FA, the shared variance between the connections weighted with these measures is limited.

**Figure 1.**
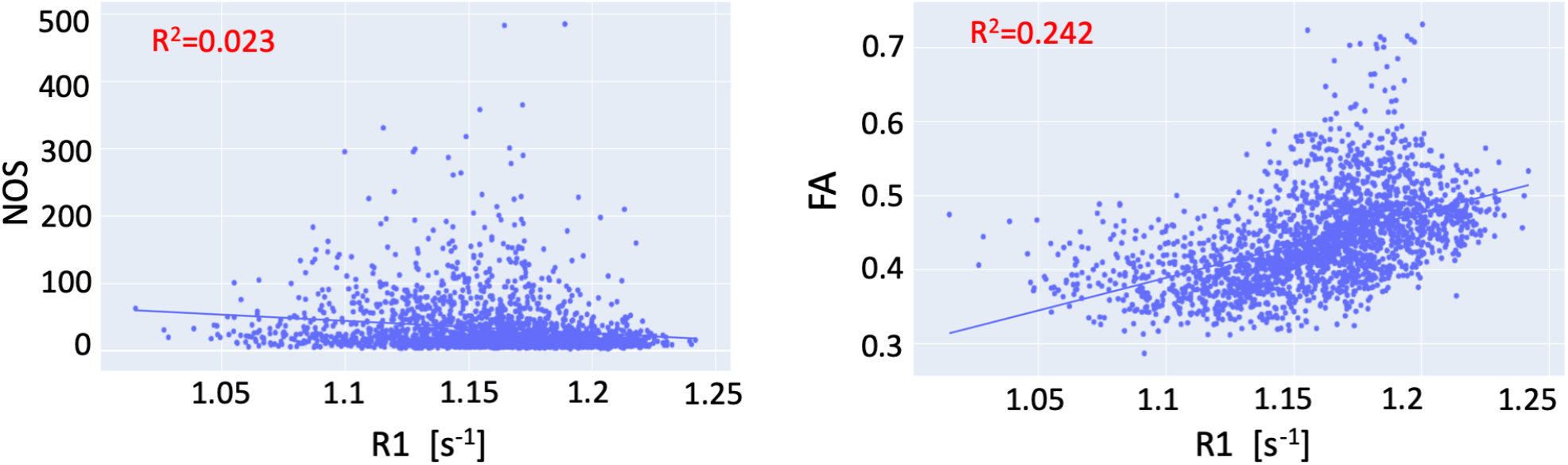
Relationship between the connection weights in the R1-weighted and FA-weighted connectome (left) and R1-weighted and NOS-weighted (right)

Then, we looked at the strength distribution and weighted average for the NOS and R1 weighted connectomes. The strength distribution of the NOS-weighted connectome is heavy-tailed (Figure 2). Among the nodes with the highest strength were the superior frontal gyrus, lateral occipital, pre- and postcentral gyrus. (Table 1 in the supplementary materials).

**Figure 2.**
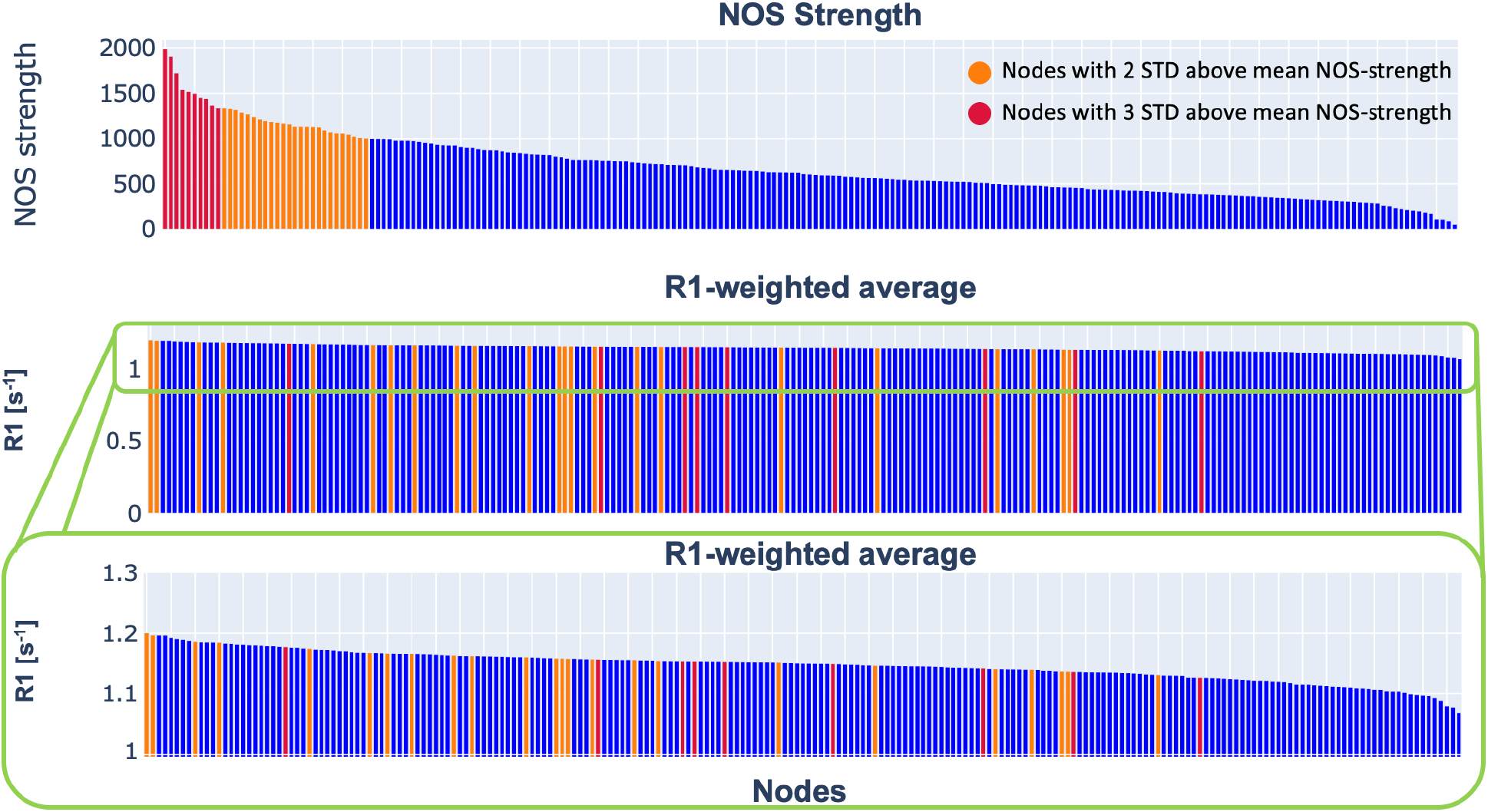
Nos strength and R1-weighted average distribution of the group NOS- and R1-weighted connectome. The plot in the middle shows the R1-weighted average distribution in the original range. However, since the trend of the R1-weighted average distribution is not visible in the original range, we selected a range that makes the trend more visible (bottom plot). In orange are highlighted the nodes that are two standard deviations above the mean NOS-strength, while in red are highlighted the nodes that are three standard deviations above the NOS-strength. The details about the nodes are provided in the supplementary materials.

On the other hand, the R1-weighted average distribution did not follow the same trend as the NOS strength distribution (Figure 2). This result indicates that a high number of streamlines is not associated with a higher median in the R1 distribution. Also, the hubs defined with the more conservative threshold (at least 3 standard deviations above the mean NOS strength) did not exhibit a high R1-weighted average (Figure 2).

As for the community structure (Figure 3), the selected resolution parameter was 0.8 for R1-weighted, while for the NOS-weighted connectome was 2.6. The consensus modularity for the R1-weighted connectome yielded 5 modules with average modularity score *Q(γ)* = 0.569. On the other hand, the NOS-weighted connectome yielded 11 modules with an average modularity score of *Q(γ)* = 0.44. We further explored the organization of the modules by looking at the distributions of the functional classes of the nodes provided by Yeo et al. (Yeo et al., 2011). Both the NOS and R1 modules were found to include multiple functional classes.

**Figure 3.**
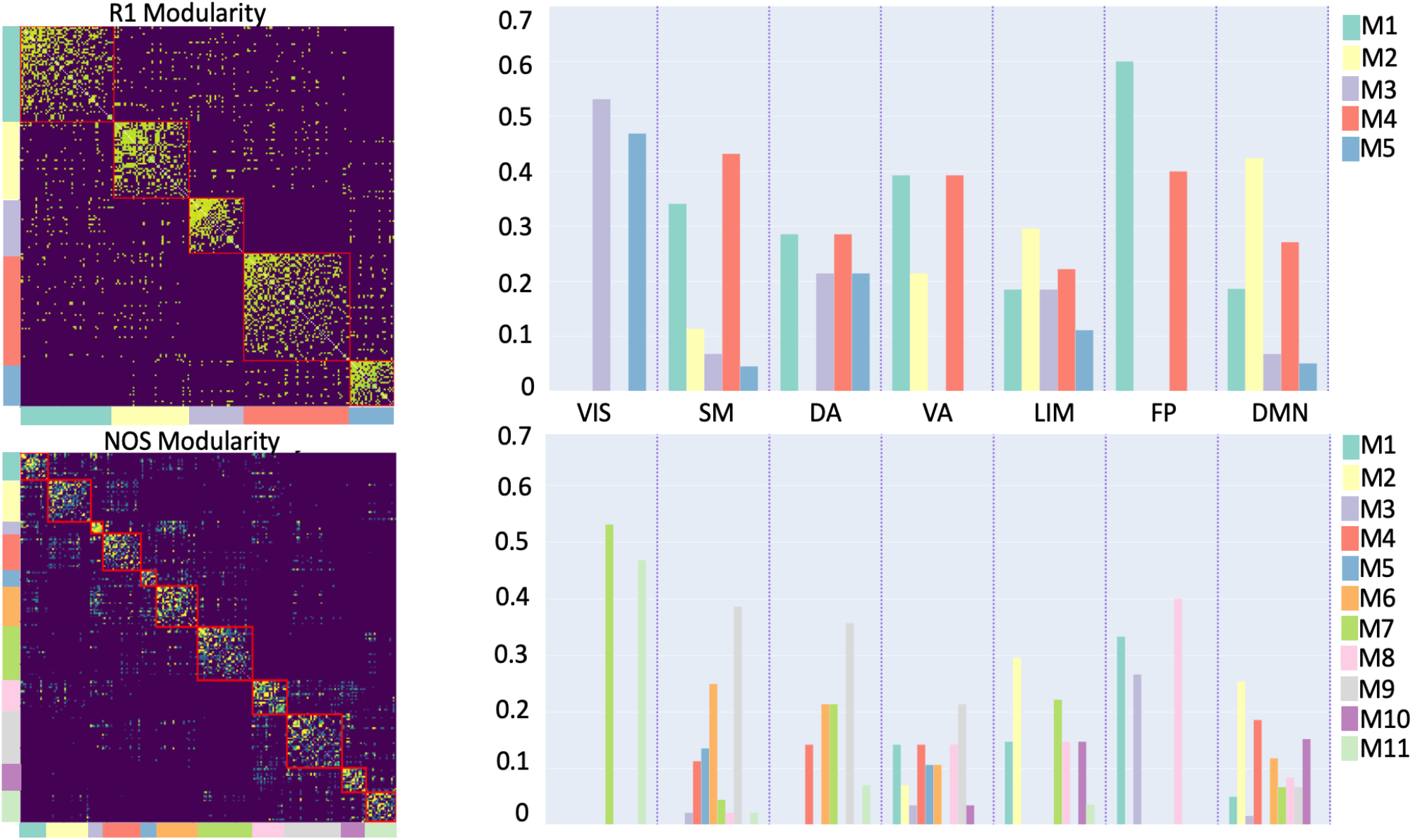
Community structure of the R1- and NOS-weighted connectomes. The bar plots represent the distributions of functional classes, given by Yeo et al. (Yeo et al., 2011), within the modules (denoted as M#) for the R1- and NOS-weighted connectomes, respectively. Yeo’s functional classes: SM (Somatomotor), VIS (Visual), VA (Ventral Attention), FP (Frontoparietal), LIM (Limbic), DA (Dorsal Attention), DMN (Default Mode Network)

The rank-based analysis (Figure 4) shows where the functional and cytoarchitectonic classes are over- and underrepresented in terms of R1-weighted average and NOS strength. For the Yeo’s functional atlas, the R1 is overrepresented, compared to NOS, in the higher-order subnetworks (transmodal) and underrepresented for function-specific subnetworks (unimodal). However, this is not the case for the cytoarchitectonic subnetworks derived using the von Economo parcellation, i.e., there was not a good distinction whether the R1 is underrepresented only for unimodal subnetworks since it was also underrepresented for the insular and the limbic transmodal subnetworks.

**Figure 4.**
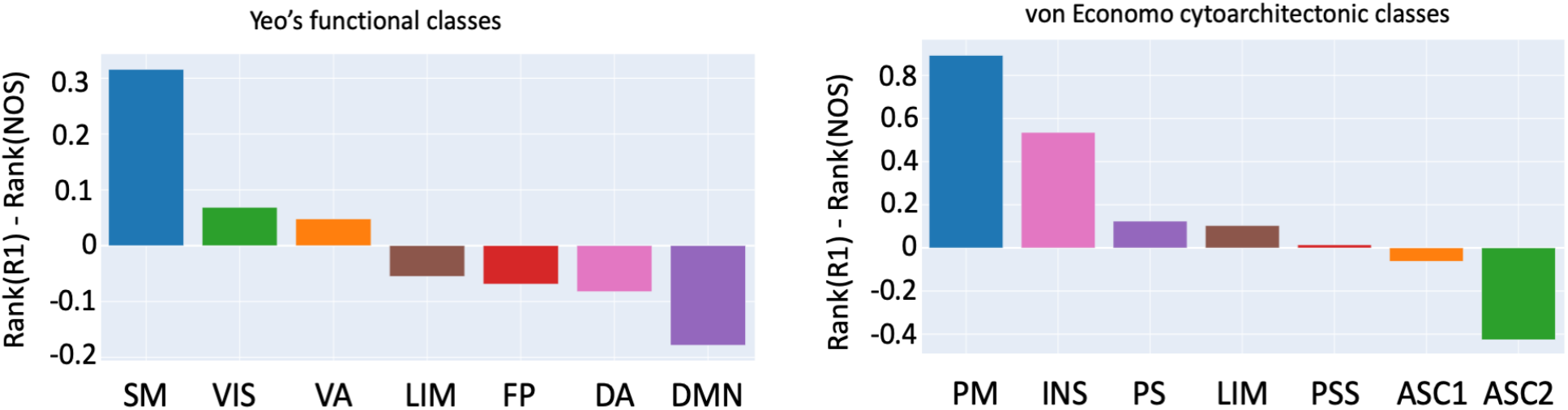
Rank-based comparison across functional and cytoarchitectonic classes. The rank for each node was calculated by its strength (for NOS)/weighted average (for R1) and then grouped using a cytoarchitectonic parcellation and functional one. Yeo’s functional classes: SM (Somatomotor), VIS (Visual), VA (Ventral Attention), FP (Frontoparietal), LIM (Limbic), DA (Dorsal Attention), DMN (Default Mode Network). Von Economo cytoarchitectonic classes: PM (primary motor), INS (insular), LIM (Limbic), PS (primary sensory), PSS (primary secondary sensory), ASC1 (association cortex), ASC2 (association cortex 2)

We repeated the same analysis on the connectomes constructed with a stricter threshold i.e. two regions are connected if there are at least 5 streamlines reconstructed between them (see Supplementary materials). The results showed that centrality measures’ distributions and rank-based analysis are consistent between the two thresholds. However, regarding the modularity, R1-based connectome yielded a different number of modules, although the community structure was still different from NOS.

## Discussion

In this study, we showed that by using a myelin sensitive measure we can complement the diffusion MRI-based connectivity and provide a different picture of the brain organization. Given the role of myelin in the rapid transfer of information in the brain, this approach can extend the current network models and potentially offer different perspectives for demyelination diseases. To characterize the complementary aspect of myelin weighted connectomes, we compared it with a connectome weighted with a common diffusion derived metric (NOS).

First, we focused on the strength distribution and compared it to the R1-weighted average. From Figure. 2 one can appreciate that the R1-weighted average distribution does not follow the same trend as the NOS strength distribution. The R1-weighted average distribution reflected a more uniform distribution. We also found that the hub regions, defined in the NOS-connectome, do not necessarily have a high R1-weighted average. Similar results have been previously reported in (Mancini et al., 2018) for a g-ratio-weighted connectome.

Second, we observed differences in the modular structure between the NOS- and R1-weighted connectomes. The number of modules was influenced by the resolution parameter, and a different number of modules was expected as the most optimal parameters were different for the two connectomes. However, what we wanted to highlight in this study was the different modular structure for the two weights, and to do this, we partitioned the network in the most appropriate way for each weight. We also explored the distribution of the functional classes within the modules and found that there was limited agreement between the functional classes and the estimated modules, i.e., the modules included multiple functional classes. This result is in agreement with results previously reported in the literature. Betzel and colleagues found that the community structures in the structural networks do not fully resemble the community structures identified in the resting-state functional brain networks (Betzel et al., 2013). In general, it has been observed that structural and functional perspectives highlight different inter-regional relationships (Goni et al., 2014; Honey et al., 2010; Suárez et al., 2020).

Regarding the rank-based analysis, we found that there was a good division of the unimodal versus transmodal functional classes. This pattern seems to follow the functional gradient observed in previous studies (Margulies et al., 2016; Vázquez-Rodríguez et al., 2019). An interesting result was that the unimodal regions exhibited more connections but in proportion a lower R1-weighted average, while the transmodal regions exhibited higher R1-weighted average but less connections. This could potentially mean that the unimodal regions need to use more connections with lower myelination for a reliable transfer of the information, whereas the transmodal regions get a reliable transfer of information through more myelinated, but fewer, connections. A recent study has shown an opposite trend in cortical gray matter (Glasser & van Essen, 2011), but our study focuses on white matter connectivity and uses a different imaging modality (R1 versus T1w/T2w).

Our results showed that differences exist between the connectome weighted with NOS and the one weighted with R1 in terms of the distribution of their weights, as well as in the modular organization. Therefore, the R1-weighted connectome can be used to study the relationship between white matter myeloarchitecture and function, given that the rank-based analysis showed an agreement in subdivision of the regions in unimodal and transmodal functional subnetworks.

The use of qMRI metrics to weight the connectome could have important implications for many applications. qMRI offers several techniques that are sensitive to myelin (Laule et al., 2007; Petiet et al., 2019), such as magnetization transfer, myelin water imaging, or relaxometry (for a complete list see (Heath et al., 2018)). Additionally, these techniques could be used to estimate the conduction velocity and conduction delays, and to incorporate these metrics as weights in the connectome. This would potentially result in a more complete model of the structural connectome and may provide a more comprehensive understanding of how the structure shapes the function. In this direction, Berman and colleagues calculated the conduction delay among the fibers in the corpus callosum using MRI-derived g-ratio (Berman et al., 2019). However, to calculate the conduction velocities and delays, in addition to the information about myelin, one would also need information about the axonal diameter and potentially about other microstructural properties not accessible from MRI (Drakesmith et al., 2019). In the work of Drakesmith et al (Drakesmith et al., 2019), they studied the feasibility of estimating conduction velocity in vivo using MRI microstructural measures. They performed simulations and reported that most of the variance in the estimation of the conduction velocity is explained by the axonal diameter and the g-ratio. Using both myelin and axonal diameter data would allow one to characterize more thoroughly brain pathologies such as multiple sclerosis, Parkinson’s or Alzheimer’s disease, more thoroughly. However, axonal diameter can be accurately measured only with high gradients (300 mT/m) (Veraart et al., 2020) and is therefore not a measure that one can have on a clinical scanner yet. Additionally, even at such high gradients, the MRI-derived axonal measure is not sensitive to small axons (1μm or lower) (D. K. Jones et al., 2018), so there are still challenges that need to be tackled in order to compute a robust estimate of the conduction velocity or delay.

There are a few methodological aspects of this work that are worth mentioning. The first is the choice of quantitative MRI metrics to weight the connectome. As mentioned before, the structural connectome is often weighted using diffusion derived metrics such as NOS and FA. For NOS, this stems from the assumption that streamline count is a proxy of microstructural fiber count, i.e., the greater the number of streamlines, the higher the connectivity between regions. This has been shown to be questionable, however, as results are influenced by the tractography algorithms and the choice of tracking parameters. FA, moreover, is traditionally considered to be an indirect measure of white matter integrity and, as such, is routinely employed as an alternative weight in structural connectomes (Martijn P Van Den Heuvel et al., 2019) but is not a direct measure of myelin since the diffusion generally is blind to myelin. The myelin has a short T2 relaxation (Heath et al., 2018), therefore it cannot be captured by the long echo times required for diffusion acquisition. To mitigate the lack of specificity of FA and NOS to the brain microstructure, we decided to use R1 to weight the connectome. The relaxation rate, R1, has been repeatedly shown to correlate highly with myelin content (Lee et al., 2012; Lutti et al., 2014). Also, the MP2RAGE sequence, which was used to acquire the R1 maps, is a stock, relatively short protocol with open-source processing, which makes it suitable for a wide clinical application. There are several studies that demonstrated the usefulness of complementing the tractography with longitudinal relaxation time. For instance, (De Santis et al., 2014) showed that to compare two groups i.e. to detect differences between groups, the longitudinal relaxation time (T1), which is just an inverse of R1, requires a smaller sample size compared to the diffusion derived metrics. Another study (De Santis et al., 2016) demonstrated that it is possible to measure tract-specific T1 relaxation, potentially leading to fiber-specific myelin metrics and more thorough network models.

Another aspect is that here we weighted the connectome using the median rather than the standard approach of taking the mean along the bundle of reconstructed streamlines. This is due to the fact that the median is more robust against outliers and does not rely on the normality assumption for the R1 distribution along a fiber bundle. Relying on one measure per bundle instead averaging a measure across streamlines also avoids biasing the results towards NOS.

Furthermore, we should also mention the choice of network measures that were investigated. The more canonical graph measures such as clustering coefficient and path length were not calculated for this study as they assume that the weight of a connection is proportional to information transfer or a related property. This is not necessarily the case for an absolute measure of myelin. For this reason, we concentrated our analysis on the centrality measures and modular structure of the connectomes.

One limitation in this study, that also exists in many connectomics studies, is the use of diffusion MRI to reconstruct the connectome. In a recent study (Maier-Hein et al., 2017), it has been shown that tractography algorithms produce a large amount of false positive streamlines which further bias the reconstructed connectome. This limitation is particularly evident for modularity: the different results obtained for R1 using two different thresholds may imply that including spurious streamlines deeply affects the R1 weight distribution and therefore the estimated modular structure. To tackle these thresholding issues, new algorithms have recently been proposed (Schiavi et al., 2020; Smith et al., 2015) that aim to reduce the number of false positive streamlines by using microstructural and anatomical priors. Future studies need to clarify how such methods could be applied in studies that combine tractography with complementary measures

In conclusion, the R1-weighted connectome complements the structural connectome derived from dMRI and could provide new biomarkers for many pathologies that affect the brain. Further validation of this approach is required, e.g. by studying demyelinating diseases.

## Data and code availability

The code and data, to reproduce the results, are available on GitHub (https://github.com/TommyBoshkovski/The_R1-weighted_connectome).

## Acknowledgement

We thank the ICEBERG study group and particularly Marie Vidailhet, MD (Pitié-Salpêtrière Hospital, Paris, Principal investigator), Jean-Christophe Corvol, MD, PhD (Paris Brain Institute, Paris, clinical and genetic data), Isabelle Arnulf, MD, PhD (Pitié-Salpêtrière Hospital, Paris, clinical and sleep data), Rahul Gaurav, MS, (Pitié-Salpêtrière Hospital, Paris, data analysis), Nadya Pyatigorskaya, MD, PhD, (Pitié-Salpêtrière Hospital, Paris, data analysis); for their help in collecting data. The research leading to these results has received funding from the programs “Investissements d’Avenir” ANR-10-IAIHU-06 and ANR-11-INBS-0006, the EDF Foundation, the Fondation Thérèse and René Planiol, Unrestricted support for Research on Parkinson’s disease from Energipole (M. Mallart) and Société Française de Médecine Esthétique (M. Legrand), Montreal Heart Institute Foundation (NS), Canadian Open Neuroscience Platform (Brain Canada PSG) (NS), Quebec Bio-Imaging Network (NS, 8436-0501), Natural Science and Engineering Research Council of Canada (NS, 2016-06774) Fonds de Recherche du Québec (NS, FRSQ 36759 and FRSQ 35250). MM was funded by the Wellcome Trust through a Sir Henry Wellcome Postdoctoral Fellowship [213722/Z/18/Z].

## Supplementary materials

In the supplementary materials we provided additional results using also the FA to weight the connectome. We also reported the outcomes obtained using a more stringent threshold in the connectivity matrices.

