## Supplementary Materials for "The R1-weighted connectome: complementing brain networks with a myelin-sensitive measure"

#### S1. Strength and weighted-average distributions

| | NOS Strength | R1-weighted average | FA-weighted average |
|---|---|---|---|
| caudalmiddlefrontal_3_R | 1966 | 1.15455223 | 0.45481415 |
| superiorfrontal_8_L | 1955.5 | 1.15489756 | 0.5364118 |
| superiorfrontal_5_R | 1725.5 | 1.15657259 | 0.48931146 |
| lateraloccipital_4_R | 1558 | 1.17902009 | 0.37937757 |
| caudalmiddlefrontal_3_L | 1510.5 | 1.1425173 | 0.45794132 |
| precentral_6_R | 1502 | 1.12808163 | 0.47504907 |
| lateraloccipital_1_L | 1484.5 | 1.15372432 | 0.4250611 |
| postcentral_5_R | 1447 | 1.13491831 | 0.44638249 |
| caudalmiddlefrontal_1_L | 1360.5 | 1.15249975 | 0.44708179 |
| superiorfrontal_4_R | 1357.5 | 1.14975925 | 0.44246636 |
| lateraloccipital_2_R | 1330 | 1.18572172 | 0.40010313 |
| superiorparietal_7_R | 1329.5 | 1.18490026 | 0.46868563 |
| superiorfrontal_6_L | 1321 | 1.13927463 | 0.43705957 |
| superiorfrontal_3_R | 1292.5 | 1.19858478 | 0.45097488 |
| inferiorparietal_6_R | 1292.5 | 1.15621096 | 0.42422677 |
| parsopercularis_2_L | 1249.5 | 1.15720478 | 0.44614924 |
| paracentral_2_L | 1241 | 1.13771093 | 0.44208708 |
| rostralmiddlefrontal_2_R | 1214.5 | 1.16733895 | 0.43347641 |
| superiorfrontal_6_R | 1193.5 | 1.15941077 | 0.53443044 |
| precentral_4_R | 1193.5 | 1.13806138 | 0.44677296 |
| superiorparietal_1_R | 1178.5 | 1.16853382 | 0.43314363 |

| | | | |
|---|---|---|---|
| superiorfrontal_7_L | 1159 | 1.15768024 | 0.49319574 |
| rostralmiddlefrontal_1_R | 1151 | 1.14500352 | 0.41263682 |
| cuneus_1_L | 1135.5 | 1.13002028 | 0.41040971 |
| precentral_7_L | 1124 | 1.15641855 | 0.43493235 |
| inferiorparietal_4_L | 1111 | 1.16574541 | 0.43595524 |
| lateraloccipital_4_L | 1095 | 1.16091219 | 0.40645647 |
| middletemporal_1_L | 1094 | 1.17478661 | 0.45568292 |
| middletemporal_1_R | 1077.5 | 1.19980463 | 0.43495825 |
| superiorfrontal_3_L | 1075 | 1.16314845 | 0.45295976 |
| superiorfrontal_9_L | 1071.5 | 1.13961489 | 0.47527627 |
| rostralmiddlefrontal_3_R | 1032 | 1.15389897 | 0.40840144 |
| lateraloccipital_5_R | 1023.5 | 1.19707436 | 0.40468686 |
| inferiorparietal_1_L | 1017 | 1.1632152 | 0.42050431 |
| precentral_5_R | 998 | 1.12466084 | 0.44015975 |
| superiorparietal_2_L | 995.5 | 1.1501535 | 0.42405723 |
| parstriangularis_1_L | 994 | 1.15047862 | 0.41179135 |
| inferiorparietal_5_L | 988 | 1.16492729 | 0.43319758 |
| isthmuscingulate_1_L | 985.5 | 1.15111918 | 0.46342115 |
| superiorparietal_7_L | 983.5 | 1.15835931 | 0.46232216 |
| precentral_3_L | 982.5 | 1.10873801 | 0.45172339 |
| superiorfrontal_7_R | 981 | 1.14491754 | 0.50266414 |
| supramarginal_5_L | 975.5 | 1.15672519 | 0.42411832 |
| parsopercularis_2_R | 971 | 1.15793737 | 0.44405594 |
| postcentral_3_R | 961.5 | 1.14229538 | 0.41660898 |

| | | | |
|---|---|---|---|
| rostralmiddlefrontal_2_L | 960 | 1.15095653 | 0.42127552 |
| superiorfrontal_2_R | 957 | 1.15379185 | 0.45018773 |
| paracentral_2_R | 923.5 | 1.13875271 | 0.4811363 |
| superiorfrontal_1_R | 923 | 1.15586281 | 0.44184927 |
| rostralmiddlefrontal_4 _L | 908.5 | 1.16348021 | 0.39311237 |
| superiorfrontal_5_L | 905 | 1.15523623 | 0.45805608 |
| superiorparietal_5_L | 889.5 | 1.16252581 | 0.43204665 |
| superiorfrontal_8_R | 887.5 | 1.14141001 | 0.44379638 |
| caudalmiddlefrontal_1_R | 883 | 1.14609962 | 0.44162341 |
| pericalcarine_1_L | 871 | 1.11404583 | 0.39693474 |
| superiorparietal_4_R | 866.5 | 1.18072143 | 0.42485591 |
| lateraloccipital_1_R | 852 | 1.15844078 | 0.42304646 |
| lateraloccipital_5_L | 849 | 1.13991544 | 0.37970863 |
| superiorfrontal_1_L | 839.5 | 1.17374917 | 0.45768595 |
| supramarginal_4_L | 839 | 1.16224559 | 0.4302635 |
| inferiorparietal_2_L | 832 | 1.16368641 | 0.42498126 |
| inferiortemporal_4_L | 818.5 | 1.15592484 | 0.4324534 |
| precentral_4_L | 815.5 | 1.10742643 | 0.43777023 |
| cuneus_2_R | 815 | 1.15082102 | 0.44632162 |
| rostralmiddlefrontal_5 _L | 807.5 | 1.17197196 | 0.39257267 |
| rostralmiddlefrontal_1_L | 792.5 | 1.16495753 | 0.42939455 |
| insula_3_L | 790 | 1.10283158 | 0.4169084 |
| isthmuscingulate_1_R | 788.5 | 1.16147066 | 0.45335847 |
| inferiortemporal_4_R | 784.5 | 1.19627428 | 0.421529 |

| | | | |
|---|---|---|---|
| parstriangularis_2_R | 776 | 1.15943345 | 0.41145212 |
| precentral_8_L | 768 | 1.13441633 | 0.42563866 |
| superiortemporal_5_R | 767 | 1.14868755 | 0.44976691 |
| superiorparietal_4_L | 763 | 1.16261628 | 0.44885281 |
| inferiorparietal_2_R | 761 | 1.18345119 | 0.41581574 |
| insula_2_R | 760.5 | 1.11639998 | 0.42300816 |
| precentral_2_R | 754.5 | 1.16312108 | 0.44005198 |
| rostralanteriorcingulate_1_L | 745.5 | 1.1462204 | 0.42977588 |
| supramarginal_2_R | 739.5 | 1.19077522 | 0.4249869 |
| precuneus_5_R | 737 | 1.16825677 | 0.46423564 |
| inferiorparietal_1_R | 736.5 | 1.18330042 | 0.43022905 |
| inferiorparietal_3_R | 730.5 | 1.19412587 | 0.43864252 |
| supramarginal_3_R | 724 | 1.17614717 | 0.40946224 |
| posteriorcingulate_1_L | 721.5 | 1.14034405 | 0.44099281 |
| medialorbitofrontal_2_L | 704.5 | 1.12784405 | 0.41264181 |
| precentral_5_L | 699.5 | 1.18694673 | 0.45829026 |
| inferiorparietal_4_R | 699.5 | 1.13859579 | 0.42150329 |
| supramarginal_1_R | 697 | 1.16800185 | 0.42261709 |
| parsopercularis_1_L | 695 | 1.15456888 | 0.4239968 |
| precentral_2_L | 680.5 | 1.11527065 | 0.45777004 |
| rostralanteriorcingulate_1_R | 680 | 1.13511194 | 0.4101604 |
| superiorparietal_3_L | 678 | 1.14502659 | 0.40302768 |
| paracentral_1_L | 674 | 1.13882637 | 0.48018474 |
| insula_3_R | 668.5 | 1.08889984 | 0.39622527 |

| | | | |
|---|---|---|---|
| inferiorparietal_5_R | 663 | 1.17867113 | 0.39738851 |
| supramarginal_2_L | 660.5 | 1.15205014 | 0.40175146 |
| rostralmiddlefrontal_4 _R | 660 | 1.15286239 | 0.3900672 |
| superiorfrontal_2_L | 658.5 | 1.1514495 | 0.44912618 |
| postcentral_5_L | 649.5 | 1.1855542 | 0.38959689 |
| fusiform_3_R | 649.5 | 1.1262997 | 0.38848968 |
| caudalmiddlefrontal_2_L | 647.5 | 1.15461277 | 0.42479 |
| lingual_3_R | 633.5 | 1.15286475 | 0.39020298 |
| middletemporal_3_L | 627 | 1.15709588 | 0.42177641 |
| paracentral_3_R | 619.5 | 1.14703501 | 0.41685414 |
| superiortemporal_3_L | 618.5 | 1.14588912 | 0.39757772 |
| superiorparietal_5_R | 612.5 | 1.18296281 | 0.42387247 |
| precentral_3_R | 604.5 | 1.15056573 | 0.42631798 |
| lateralorbitofrontal_1_L | 602.5 | 1.1397014 | 0.36462349 |
| rostralmiddlefrontal_3_L | 595 | 1.16592398 | 0.40353479 |
| parsopercularis_1_R | 593.5 | 1.15170139 | 0.42685304 |
| lingual_1_L | 593.5 | 1.12772303 | 0.39827912 |
| lateralorbitofrontal_2_R | 592 | 1.11875655 | 0.38789434 |
| medialorbitofrontal_1_R | 590 | 1.14103453 | 0.42591573 |
| rostralmiddlefrontal_5 _R | 586 | 1.15616708 | 0.39314514 |
| precuneus_2_L | 585 | 1.15801093 | 0.44607199 |
| lateraloccipital_3_R | 585 | 1.15698507 | 0.35949723 |
| postcentral_4_L | 584 | 1.10408324 | 0.41106983 |
| postcentral_3_L | 575.5 | 1.11841811 | 0.40030039 |

| | | | |
|---|---|---|---|
| precentral_1_L | 569.5 | 1.12317379 | 0.48183509 |
| posteriorcingulate_2_R | 564 | 1.15623144 | 0.42528901 |
| superiorfrontal_4_L | 563 | 1.15482501 | 0.45464763 |
| caudalanteriorcingulate_1_R | 562 | 1.13303653 | 0.42056307 |
| fusiform_4_L | 551.5 | 1.13483781 | 0.43284576 |
| caudalanteriorcingulate_1_L | 551.5 | 1.12405917 | 0.42149176 |
| fusiform_2_L | 551 | 1.14184332 | 0.38525191 |
| pericalcarine_1_R | 544 | 1.13457765 | 0.38293242 |
| fusiform_2_R | 537.5 | 1.17439895 | 0.37263486 |
| precentral_1_R | 534 | 1.14323869 | 0.42110269 |
| supramarginal_4_R | 531 | 1.14736172 | 0.36603971 |
| lateraloccipital_3_L | 529 | 1.14020491 | 0.3774293 |
| superiorparietal_3_R | 526 | 1.18642675 | 0.43409549 |
| precuneus_4_R | 524 | 1.17751237 | 0.4590447 |
| lingual_1_R | 521.5 | 1.15579823 | 0.37019401 |
| postcentral_7_L | 520.5 | 1.12540779 | 0.37327419 |
| fusiform_1_R | 520 | 1.1691619 | 0.34769522 |
| postcentral_1_L | 517.5 | 1.141686 | 0.47087571 |
| fusiform_1_L | 515.5 | 1.13122178 | 0.34963292 |
| superiorparietal_6_R | 513.5 | 1.1832551 | 0.40556838 |
| parsorbitalis_1_R | 512.5 | 1.14404038 | 0.38672418 |
| precuneus_3_L | 506.5 | 1.16370884 | 0.46374493 |
| superiortemporal_5_L | 501.5 | 1.11135013 | 0.4596908 |
| fusiform_4_R | 500 | 1.16894616 | 0.40645676 |

| | | | |
|---|---|---|---|
| middletemporal_2_L | 495.5 | 1.17035314 | 0.45461105 |
| middletemporal_4_R | 489 | 1.16885257 | 0.41759972 |
| rostralmiddlefrontal_6_L | 485 | 1.17903179 | 0.40397749 |
| cuneus_1_R | 481.5 | 1.15410335 | 0.38543692 |
| inferiorparietal_3_L | 481 | 1.14867184 | 0.40587203 |
| lateralorbitofrontal_2_L | 478 | 1.12837932 | 0.38516694 |
| medialorbitofrontal_1_L | 468 | 1.15228688 | 0.42814361 |
| superiorparietal_1_L | 466.5 | 1.16181001 | 0.51192899 |
| inferiortemporal_3_L | 463.5 | 1.12694042 | 0.41518689 |
| paracentral_1_R | 452 | 1.14627292 | 0.46229125 |
| rostralmiddlefrontal_6_R | 451 | 1.15557603 | 0.38230397 |
| superiortemporal_1_L | 450.5 | 1.15113884 | 0.40836503 |
| precuneus_1_L | 448.5 | 1.16877606 | 0.50295654 |
| precuneus_5_L | 442.5 | 1.13045906 | 0.39570302 |
| postcentral_2_R | 442 | 1.14496349 | 0.39419567 |
| lateralorbitofrontal_4_L | 439 | 1.16620957 | 0.41169347 |
| superiortemporal_3_R | 432 | 1.15778882 | 0.38670591 |
| superiorparietal_6_L | 429.5 | 1.13243551 | 0.38101755 |
| lateraloccipital_2_L | 426 | 1.11668654 | 0.37404023 |
| posteriorcingulate_2_L | 425 | 1.12638331 | 0.39814242 |
| posteriorcingulate_1_R | 424.5 | 1.13274037 | 0.41780136 |
| lingual_2_R | 418 | 1.14083757 | 0.34839059 |
| fusiform_3_L | 417 | 1.13187888 | 0.37801114 |
| inferiortemporal_1_R | 412.5 | 1.15713411 | 0.40578383 |

| | | | |
|---|---|---|---|
| supramarginal_3_L | 410.5 | 1.15168448 | 0.4131758 |
| superiortemporal_4_L | 410.5 | 1.1323127 | 0.4079409 |
| superiortemporal_2_R | 407.5 | 1.17260683 | 0.38742364 |
| superiorparietal_2_R | 398 | 1.18109749 | 0.52135756 |
| insula_1_L | 393 | 1.10346004 | 0.3695119 |
| postcentral_1_R | 379 | 1.14983243 | 0.43441182 |
| middletemporal_4_L | 379 | 1.13664358 | 0.37230518 |
| bankssts_1_R | 375 | 1.17994353 | 0.392723 |
| precentral_6_L | 373.5 | 1.10567908 | 0.37237818 |
| lingual_2_L | 369.5 | 1.12052779 | 0.36558653 |
| caudalmiddlefrontal_2_R | 364 | 1.14904611 | 0.4179389 |
| insula_1_R | 363.5 | 1.13455426 | 0.37337997 |
| precuneus_3_R | 363 | 1.16865013 | 0.40939818 |
| precuneus_4_L | 362.5 | 1.15330462 | 0.40953028 |
| superiortemporal_1_R | 360.5 | 1.16366432 | 0.38950814 |
| lateralorbitofrontal_3_L | 357.5 | 1.14780076 | 0.3412609 |
| lingual_4_L | 351 | 1.11990501 | 0.35407521 |
| precuneus_2_R | 346 | 1.16974181 | 0.4330711 |
| supramarginal_1_L | 342.5 | 1.12020763 | 0.354774 |
| insula_2_L | 341 | 1.10332633 | 0.41775601 |
| frontalpole_1_R | 340.5 | 1.17891993 | 0.47641935 |
| parsorbitalis_1_L | 340 | 1.15781349 | 0.37857484 |
| bankssts_1_L | 338 | 1.14517744 | 0.39097622 |
| middletemporal_2_R | 336.5 | 1.18898979 | 0.41242934 |

| | | | |
|---|---|---|---|
| middletemporal_3_R | 332.5 | 1.17653929 | 0.40450314 |
| inferiortemporal_2_L | 332 | 1.11673579 | 0.40807257 |
| medialorbitofrontal_2_R | 325 | 1.12000391 | 0.40876957 |
| postcentral_6_L | 319 | 1.12089407 | 0.38173278 |
| inferiortemporal_1_L | 316.5 | 1.1110917 | 0.3982293 |
| lingual_3_L | 314.5 | 1.10730392 | 0.37253977 |
| superiortemporal_2_L | 311.5 | 1.14632003 | 0.39161942 |
| precuneus_1_R | 309.5 | 1.14464919 | 0.39955046 |
| pericalcarine_2_R | 309 | 1.10683551 | 0.36736579 |
| bankssts_2_L | 303.5 | 1.15361921 | 0.41426406 |
| inferiortemporal_2_R | 303 | 1.14625646 | 0.39568039 |
| postcentral_2_L | 297.5 | 1.09886111 | 0.44066292 |
| lateralorbitofrontal_1_R | 291 | 1.08972515 | 0.39367714 |
| lateralorbitofrontal_3_R | 285 | 1.11050945 | 0.37323283 |
| parahippocampal_1_R | 284.5 | 1.13200874 | 0.37900699 |
| transversetemporal_1_L | 276 | 1.1177173 | 0.3515679 |
| parahippocampal_1_L | 274.5 | 1.09368955 | 0.3970375 |
| frontalpole_1_L | 269.5 | 1.18547491 | 0.47789238 |
| inferiortemporal_3_R | 235 | 1.15869384 | 0.39527496 |
| lateralorbitofrontal_4_R | 224.5 | 1.12652473 | 0.39936488 |
| postcentral_4_R | 221.5 | 1.14408016 | 0.39030834 |
| transversetemporal_1_R | 216 | 1.13834961 | 0.3555768 |
| parstriangularis_1_R | 190 | 1.09673742 | 0.38169027 |
| insula_4_L | 185.5 | 1.07750943 | 0.3689944 |

| | | | |
|---|---|---|---|
| medialorbitofrontal_3_R | 147.5 | 1.08856092 | 0.3654355 |
| temporalpole_1_L | 122 | 1.09801082 | 0.40922319 |
| superiortemporal_4_R | 101 | 1.055135 | 0.374883 |
| entorhinal_1_L | 82.5 | 1.05744297 | 0.40704466 |
| entorhinal_1_R | 71.5 | 1.09199238 | 0.40333935 |
| temporalpole_1_R | 49 | 1.10473714 | 0.361872 |

### S2. Additional Analyses

*Comparison with FA-weighted networks*

As an additional comparison, we built a FA-weighted connectome. The procedure for obtaining the FA-weighted network is the same as for the R1-weighted connectome: we assigned to each connection the median FA value along the bundle of streamlines connecting pairs of regions.

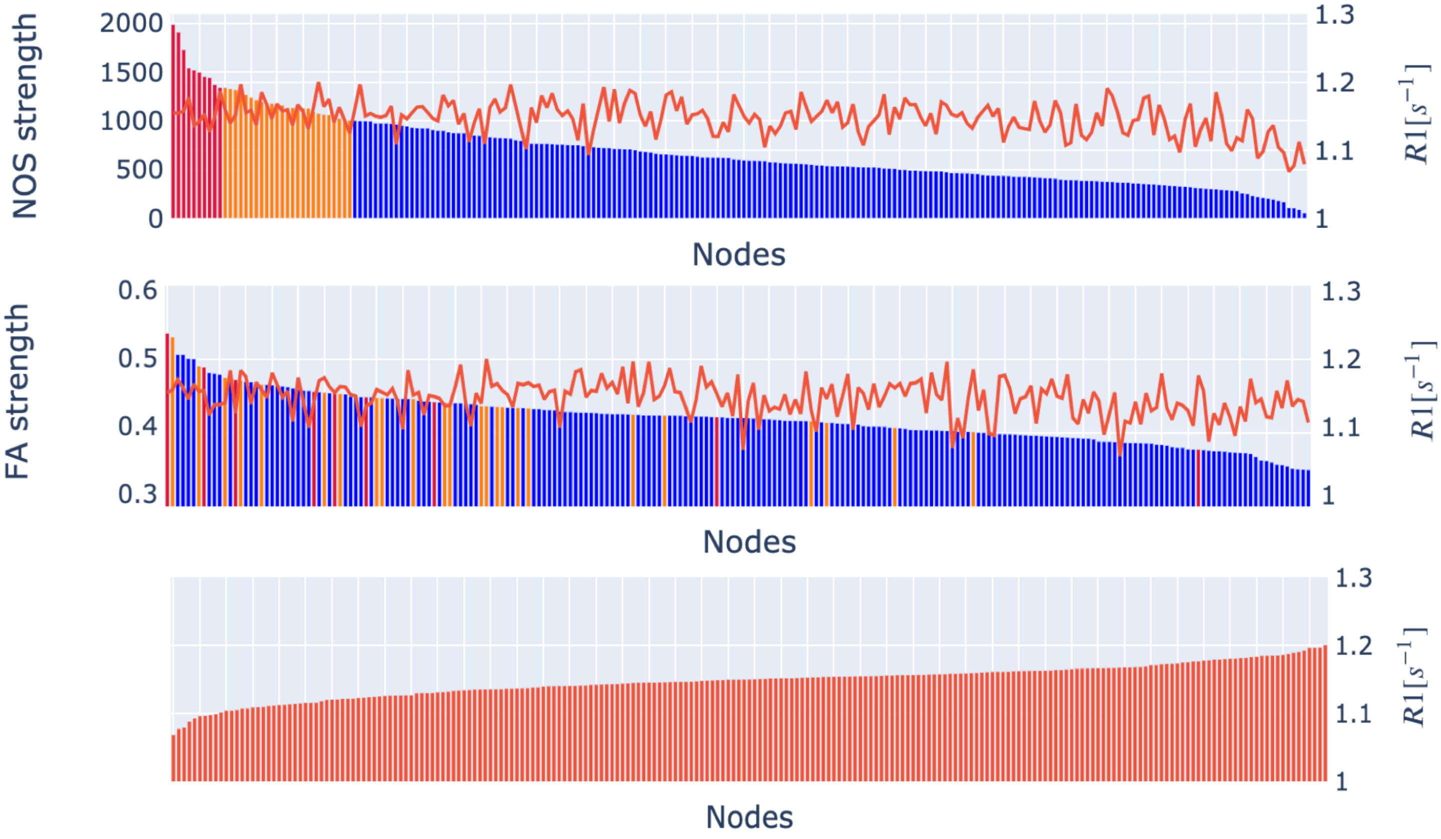


**Figure S1. Strength and weighted average distribution of the group NOS-, FA-, and R1-weighted connectome. In orange are highlighted the nodes that are two standard deviations above the mean NOS-strength, while in red are highlighted the nodes that are three standard deviations above the NOS-strength.**

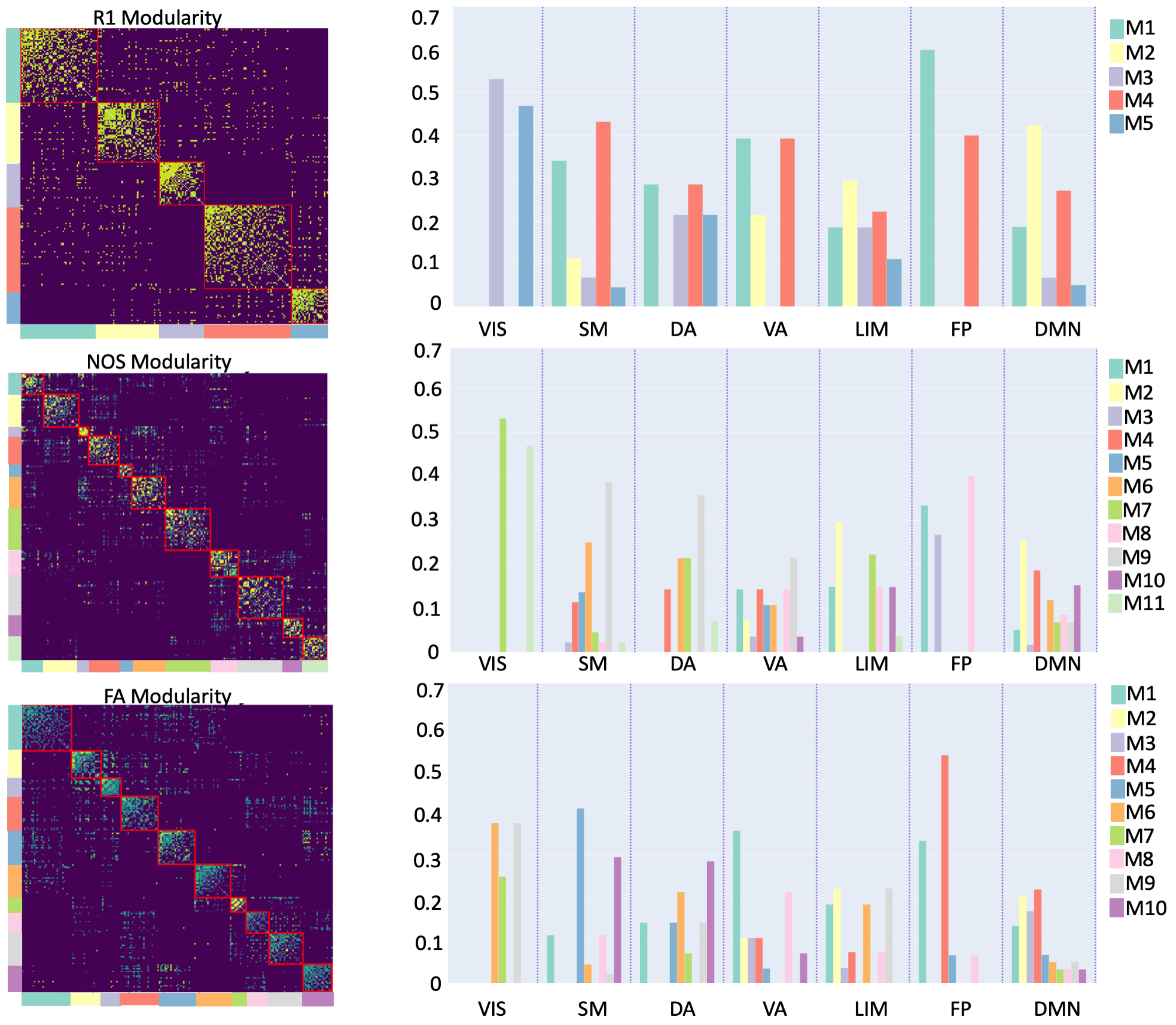


**Figure S2. Community structure of the R1-, FA-, and NOS-weighted connectomes. The bar plots represent the distributions of functional classes, given by Yeo et al., within the modules for the R1-, NOS-, and FA-weighted connectomes, respectively. Yeo's functional classes: SM (Somatomotor), VIS (Visual), VA (Ventral Attention), FP (Frontoparietal), LIM (Limbic), DA (Dorsal Attention), DMN (Default Mode Network)**

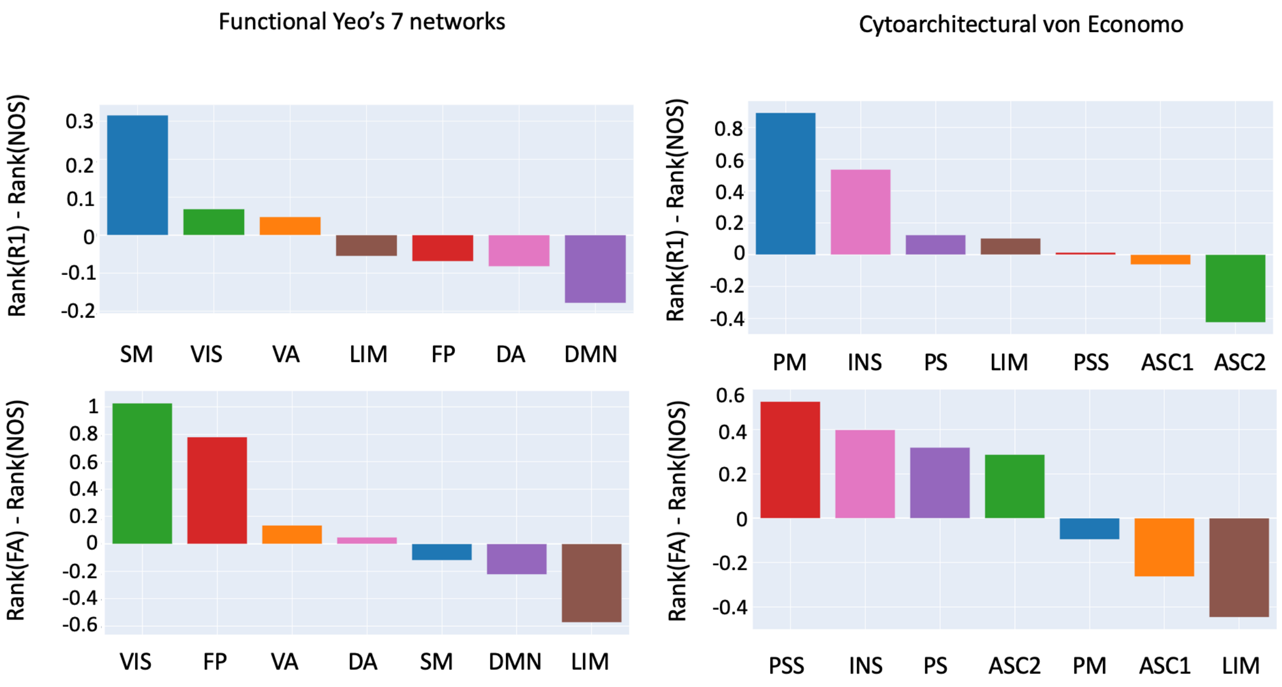


**Figure S3. Gradient of the nodes' rank. The rank for each node was calculated by its strength (for NOS)/weighted average (for R1 and FA) and then grouped using a cytoarchitectonic parcellation and functional one. Yeo's functional classes: SM (Somatomotor), VIS (Visual), VA (Ventral Attention), FP (Frontoparietal), LIM (Limbic), DA (Dorsal Attention), DMN (Default Mode Network). Von Economo cytoarchitectonic classes: PM (primary motor), INS (insular), LIM (Limbic), PS (primary sensory), PSS (primary secondary sensory), ASC1 (association cortex), ASC2 (association cortex 2)**

#### *Robustness of the analysis*

In order to check the robustness of the analyses, we built a connectome that was constructed using a more conservative threshold. Basically, we considered two nodes to be connected if and only if they are at least 5 streamlines reconstructed between them.

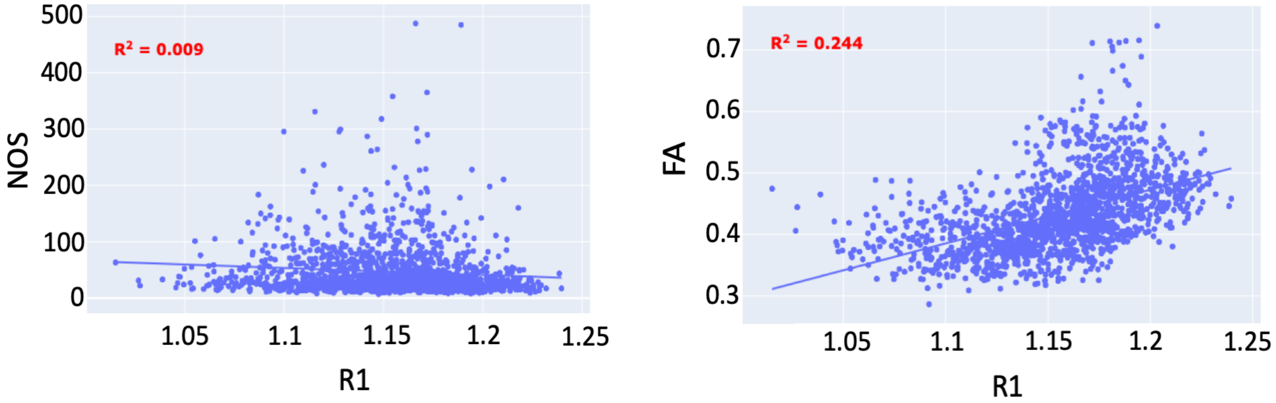


**Figure S4. Relationship between the connection weights in the R1-weighted and FA-weighted connectome (left) and R1-weighted and NOS-weighted (right )**

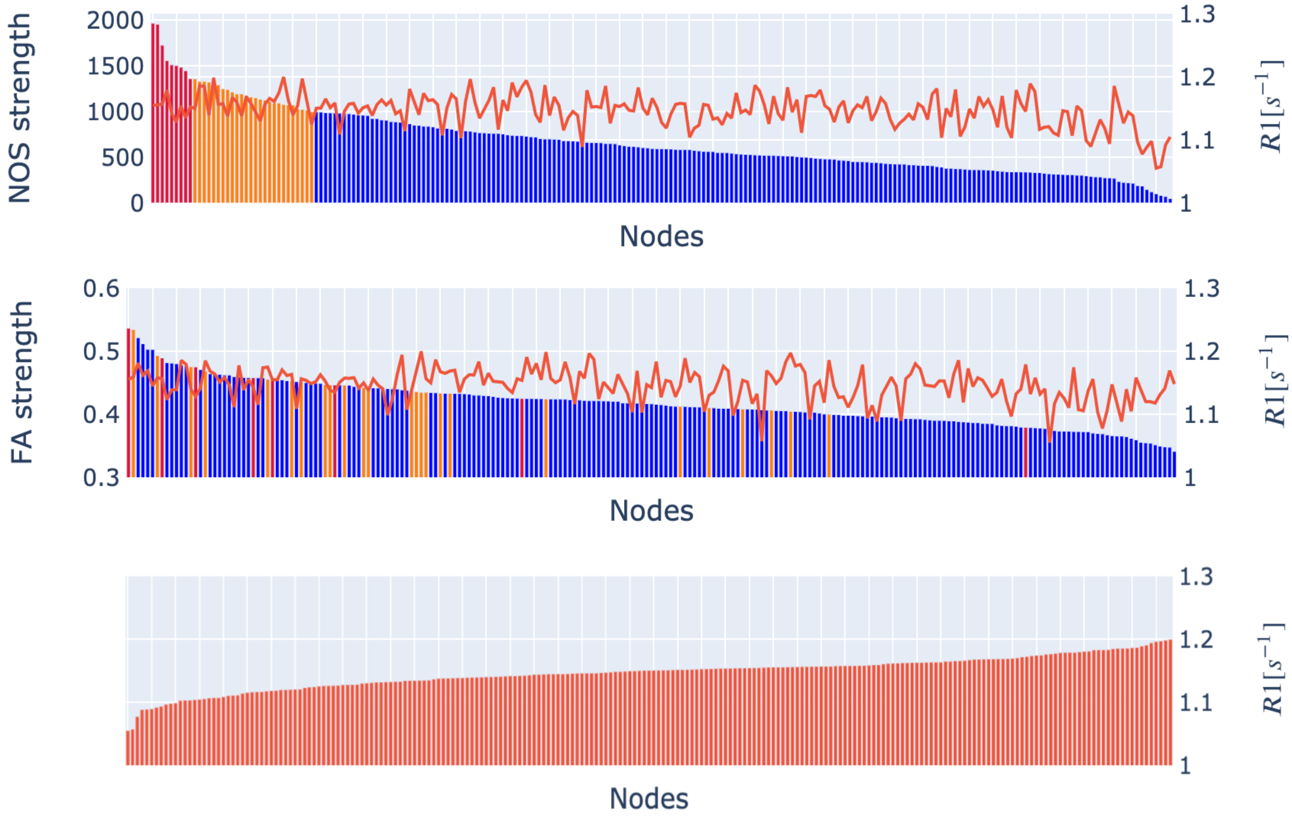


**Figure S5. Strength and weighted average distribution of the group NOS-, FA-, and R1-weighted connectome. In orange are highlighted the nodes that are two standard deviations above the mean NOS-strength, while in red are highlighted the nodes that are three standard deviations above the NOS-strength.**

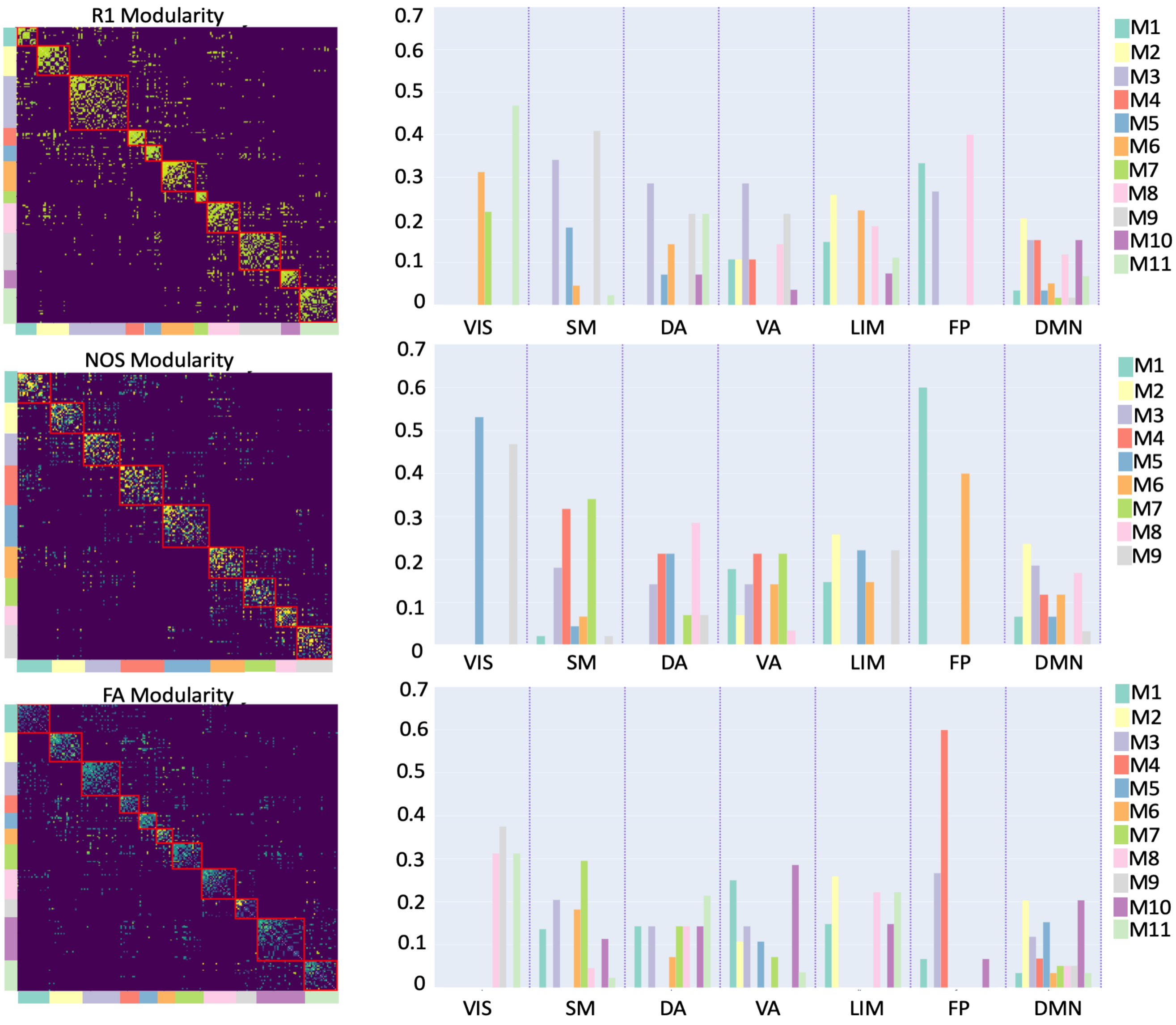


**Figure S6. Community structure of the R1-, FA-, and NOS-weighted connectomes. The selected resolution parameter was 2.8 for the R1-weighted connectome, 2.6 for the FA-weighted connectome, and 2 for the NOS-weighted connectome. The bar plots represent the distributions of functional classes, given by Yeo et al., within the modules for the R1-, NOS-, and FA-weighted connectomes, respectively. Yeo's functional classes: SM (Somatomotor), VIS (Visual), VA (Ventral Attention), FP (Frontoparietal), LIM (Limbic), DA (Dorsal Attention), DMN (Default Mode Network)**

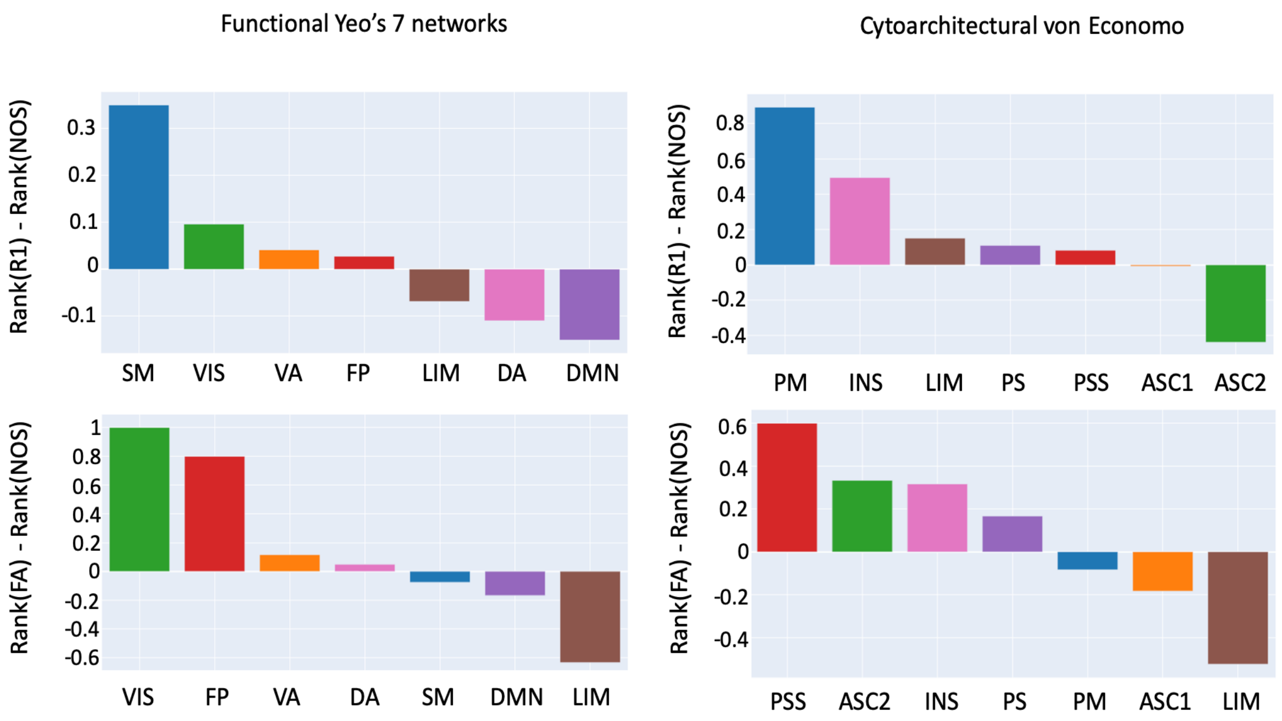


**Figure S7. Gradient of the nodes' rank. The rank for each node was calculated by its strength (for NOS)/weighted average (for R1 and FA) and then grouped using a cytoarchitectonic parcellation and functional one. Yeo's functional classes: SM (Somatomotor), VIS (Visual), VA (Ventral Attention), FP (Frontoparietal), LIM (Limbic), DA (Dorsal Attention), DMN (Default Mode Network). Von Economo cytoarchitectonic classes: PM (primary motor), INS (insular), LIM (Limbic), PS (primary sensory), PSS (primary secondary sensory), ASC1 (association cortex), ASC2 (association cortex 2)**
